# ACE2 glycans preferentially interact with the RBD of SARS-CoV-2 over SARS-CoV

**DOI:** 10.1101/2021.04.29.442038

**Authors:** Atanu Acharya, Diane L. Lynch, Anna Pavlova, Yui Tik Pang, James C. Gumbart

**Affiliations:** School of Physics, Georgia Institute of Technology, Atlanta, GA, USA

## Abstract

We report a distinct difference in the interactions of the glycans of the host-cell receptor, ACE2, with SARS-CoV-2 and SARS-CoV S-protein receptor-binding domains (RBDs). Our analysis demonstrates that the ACE2 glycan at N90 may offer protection against infections of both coronaviruses, while the ACE2 glycan at N322 enhances interactions with the SARS-CoV-2 RBD. The interactions of the ACE2 glycan at N322 with SARS-CoV RBD are blocked by the presence of the RBD glycan at N357 of the SARS-CoV RBD. The absence of this glycosylation site on SARS-CoV-2 RBD may enhance its binding with ACE2.

---

Coronaviruses have been responsible for multiple infectious diseases in the last two decades with significant global impact, including most recently COVID-19 caused by the SARS-CoV-2 virus. The SARS-CoV-2 virus binds to the angiotensin-converting enzyme 2 (ACE2) using the spike glycoprotein. While ACE2 is the receptor for both SARS-CoV-2 and SARS-CoV, it binds more strongly to the SARS-CoV-2 receptor binding domain.^1–5^

The receptor-binding motif (RBM) is composed of S438 to Q506 in SARS-CoV-2 and T425 to Q492 in SARS-CoV.^6^ A total of 17 residues of SARS-CoV-2 and 16 residues of SARS-CoV were identified to be within 4Å of ACE2, all part of their respective RBMs except for K417 of SARS-CoV-2.^6,7^ Computational studies have attributed the difference in binding strength of the coronavirus receptor-binding domains (RBDs) with ACE2 to the difference in interactions between RBD and ACE2 residues.^8–10^ Furthermore, the roles of interfacial RBD residues in the initial binding process have been analyzed by free-energy perturbation (FEP) calculations and network analysis.^9,11,12^ However, these studies do not include glycans in their respective computational models. Most studies focused on the role of SARS-CoV-2 glycans in the closed-to-open transition of the spike glycoprotein and subsequent binding to ACE2,^13,14^ as well as their impact on antibody recognition.^15^ The importance of ACE2 glycans has been less examined, especially in the context of the differences between SARS-CoV-2 and SARS-CoV. Only a few studies have investigated the interactions of ACE2 glycans with SARS-CoV-2 RBD.^16–18^ Barros et al. showed that the ACE2 glycan at N53 stabilizes the RBD-ACE2 complex and the ACE2 homodimer interface for SARS-CoV-2.^18^ Cao et al. highlighted that the glycan at N90 enhances RBD-ACE2 binding using steered molecular dynamics (SMD) simulations,^19^ although heterogeneity of glycan structures was not considered. Herein, we determine the roles of all ACE2 glycans in differential RBD-ACE2 interactions between SARS-CoV-2 and SARS-CoV.

The ACE2 peptidase domain contains up to seven N-linked and one O-linked glycans at positions N53, N90, N103, N322, N432, N546, N690, and S155. RBDs of SARS-CoV-2 and SARS-CoV contain one (at N343) and two N-glycans (at N330 and N357), respectively.^14,20^ Sequence alignment shows that the N330 and N357 amino acids of SARS-CoV RBD align with N343 and N370 of SARS-CoV-2 RBD, respectively. However, N370 of SARS-CoV-2 is not glycosylated since the requisite T359 in SARS-CoV is replaced by A372 in SARS-CoV-2. We modeled two RBD-ACE2 systems for the two viruses by adding appropriate glycans at the N- and O-glycosylation sites as suggested by mass spectroscopy experiments.^14^ Since the heterogeneity of glycans at each site does not significantly impact RBD-ACE2 interactions,^16^ we selected one glycan for each site with the largest population from the mass spectroscopy experiments. Furthermore, occupancy and processing of glycan at a specific site depends on the cell type. They can also differ in the same cell type depending on environmental conditions.^21,22^ It is common practice to perform MD simulations with multiple glycosylation patterns to account for their micro-heterogeneity.^13,16^ Therefore, we repeated all simulations with a second glycosylation pattern based on the glycan occupancy initially proposed by Azadi and co-workers.^20^ Models of the attached glycans at each site are shown in Figs. S1 and S2.

Root mean square fluctuations (RMSF) analysis reveals similar fluctuations of RBD residues in SARS-CoV and SARS-CoV-2 with the largest difference observed in a loop containing SARS-CoV residues 367-387 (380-400 in SARS-CoV-2; Fig. S3a-c). Similar RMSF profiles of the RBDs were obtained in our prior study without full glycosylations.^12^ However, none of the RBD residues from this loop interact with ACE2.^6^ Furthermore, the RMSF profiles of ACE2 residues appear very similar between the two complexes with RBD (Fig. S3a-c).

The ACE2 glycan at N322 has three distinct contact regions with both coronaviruses composed of residues 369 to 378, 404, 405, 408, 437, 439, 440, 503, 504, and 508 (SARS-CoV-2 numbering; Fig. 1a,b). Details of these contacts protocol are provided in ESI and average values are tabulated in Tables S1 and S2. Together, these contact regions of the ACE2 glycan at N322 appear on either side of the five-stranded anti-parallel *β*-sheet-domain of SARS-CoV-2 RBD. Most of the SARS-CoV-2 RBD residues interacting with the glycan at N322 are distant from the RBD-ACE2 interface. Only residues in contact region #3 (439, 440, 503, 504, and 506) lie in the RBM. Furthermore, the average number of contacts of the glycan at N322 with residues 503 and 504 is low, while it interacts with RBM residues 439 and 440 more frequently (Tables S1 and S2). The poses of the glycan at N322 when interacting with SARS-CoV-2 RBDs are similar in both glycosylation schemes adopted in this study (Fig. 1c,d). Furthermore, the glycan at N322 only interacts with three charged residues, none of which are in the RBM. The highest contact for this glycan is formed in region #1, which is composed mostly of uncharged polar residues. Calculation of the average number of contacts between RBD and ACE2 (Fig. S5a-c) shows that the overall interactions remain similar between two coronaviruses and two glycan schemes, consistent with RBD-ACE2 simulations of SARS-CoV-2 with different glycans.^16^ Structural studies also found similar interactions between SARS-CoV-2 RBD and ACE2.^4,6^

**Figure 1.**
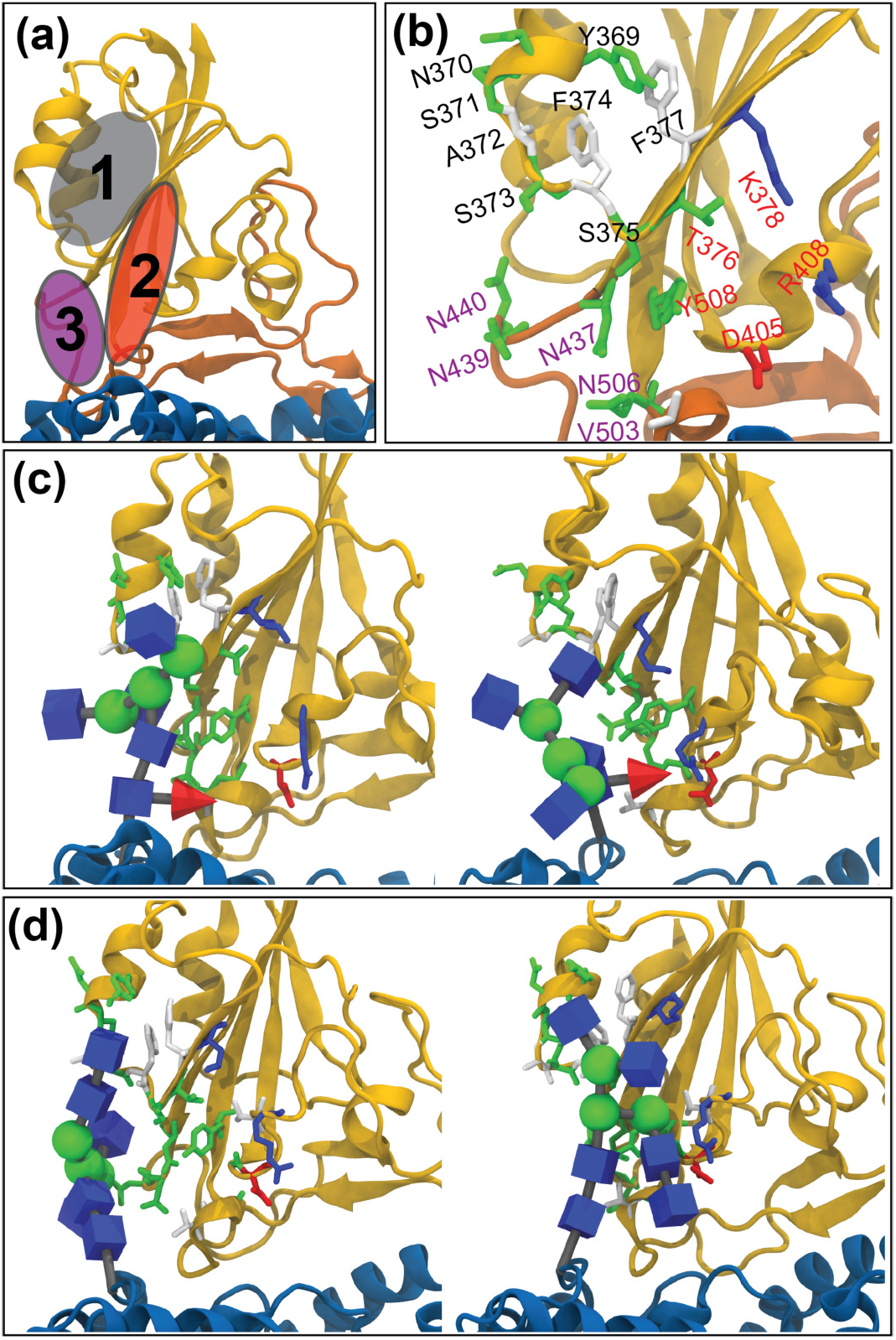
Interactions between the ACE2 glycan at N322 and SARS-CoV-2 RBD. The RBD and ACE2 are rendered in yellow and blue, respectively. (a) Three contact regions on the RBD for the glycan at N322 are marked 1, 2, and 3. RBM is highlighted in orange. (b) SARS-CoV-2 RBD residues interacting with the ACE2 glycan at N322. G404 and G504 residues are not shown. Residues in regions 1, 2, and 3 are denoted in black, red, and purple. Representative poses for the ACE2 glycan blocks at N322 interacting with SARS-CoV-2 RBD with glycosylation (c) scheme #1 and (d) scheme #2.

The glycans fluctuate significantly in MD simulation, as seen in other studies,^13,14^ and cover a significant portion of ACE2 and RBDs (Figs. 2a and S4a). We determined the average number of contacts between each ACE2 glycan and residues of coronavirus RBDs. We found that the ACE2 glycans at N90 and N322 make significant contact (Figs. 2b and S4b) with the RBDs, while the ACE2 glycans at N53 and N546 make minor contact (Fig. S6a). The ACE2 glycan at N53 plays an important role in stabilizing the homodimer interface of ACE2.^18^ We observed that independent of the glycosylation scheme, the glycan at N322 interacts more strongly with SARS-CoV-2 RBD compared to SARS-CoV. Furthermore, with glycosylation scheme #2, the glycan at N322 does not form any interaction with SARS-CoV RBD (Fig. S4b). We investigated the reasons behind striking differences in the interactions of the two coronavirus RBDs with the ACE2 glycan at N322. Sequence alignment shows that the NST(T359) sequon in SARS-CoV is replaced with NSA(A372) in SARS-CoV-2, thereby eliminating N370 as a glycosylation site in the latter coronavirus. Figs. S6b and S4c show the RBD residues covered by RBD glycans of each coronavirus. We observed that the RBD glycans at N357 and N330 of SARS-CoV block the ACE2 glycan at N322 from interacting with the RBD. In contrast, the absence of a glycan at N370 of SARS-CoV-2 RBD exposes additional sites for ACE2 glycan interaction (Figs. 2c and S4a), making this region of the RBD more susceptible to interaction with the ACE2 glycan at N322. Fig. 2c illustrates representative contact poses of relevant glycans, where the RBD glycan at N357 physically blocks the ACE2 glycan’s (at N322) contact regions. In contrast, these regions of RBD become available to stabilize the interaction with the ACE2 glycan at N322 in SARS-CoV-2. Contact calculations between SARS-CoV glycans and RBD residues reveal that the RBD glycan at N357 interacts with SARS-CoV residues 356, 364, and 366 located in the contact regions of the glycan at N322 on RBD (Figs. S6b and S4c). The RBD glycan at N343 in SARS-CoV-2 (N330 in SARS-CoV) also shows contact with some of the contact regions of the ACE2 glycan at N322. However, further investigation of the simulation trajectories reveals that the RBD glycan at N343 contacts the back of contact regions of the ACE2 glycan at N322 (Fig. 2c) and, consequently, does not block the ACE2 glycan at N322 from interacting with the RBD. Collectively, our results suggest that interaction with the regions 1 and 3 residues is crucial for binding of the ACE2 glycan at N322 to coronavirus S-protein RBDs.

**Figure 2.**
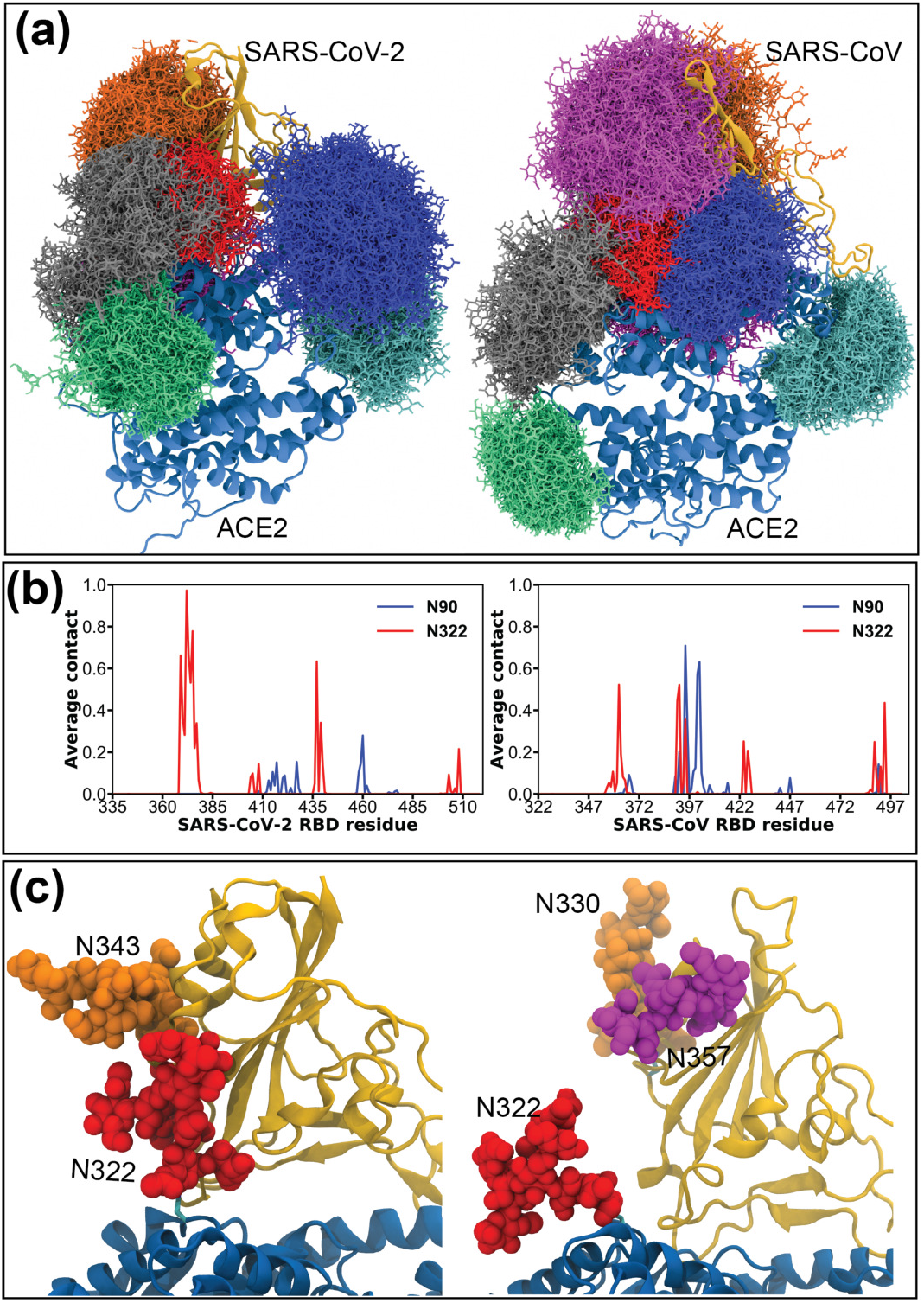
Analysis of glycan interactions. RBD and ACE2 are shown in cartoon representations in yellow and blue, respectively. (a) Comparison of glycan coverage of RBD-ACE2 complex. RBD glycan at N330 (or N343) and N357 are shown in orange and magenta colors, respectively. ACE2 glycans at N53, N90, N103, N322, N432, and N546 are shown in purple, blue, cyan, red, green, and grey colors, respectively. (b) Average number of contacts between ACE2 glycans at N90 and N322 and RBD residues of SARS-CoV-2 (left) and SARS-CoV (right) with glycosylation scheme #1 defined in Fig. S1. (c) Sample configurations of relevant glycans around RBDs of (left) SARS-CoV-2 and (right) SARS-CoV. Results from simulations of glycosylation scheme #2 are shown in Fig. S4.

Deep mutational studies revealed that mutations of most of the identified SARS-CoV-2 RBD residues modestly reduce ACE2 binding.^23^ For example, mutations of Y369-K378, D405, R408, N440, V503, G504, and Y508 of SARS-CoV-2 residues slightly reduce ACE2 binding affinity while mutations of residues N437 and N439 significantly reduce it. Only N439 and G504 SARS-CoV-2 residues are part of the RBM. Reduction in ACE2 binding with mutations of other aforementioned residues may be explained in part by reduction in the binding of the ACE2 glycan at N322. For example, most mutations of G404 and N437 in SARS-CoV-2 decrease ACE2 binding, although none of these residues are at the RBD-ACE2 interface. Therefore, an ACE2 glycan at N322 likely increases binding affinity of SARS-CoV-2 RBD with ACE2. Such an increase is absent in SARS-CoV since the RBD glycan at N357 occludes this binding site. Most mutations of T324, which eliminate the ACE2 glycan at N322, reduce binding between ACE2 and SARS-CoV-2 RBD.^24^ The absence of the RBD glycan at N370 in SARS-CoV-2 has also been suggested to increase the availability of the open spike conformation since a complex N370 glycan, when artificially introduced in SARS-CoV-2, can wrap around and tie the RBD down like a shoelace.^25^

In contrast to the ACE2 glycan at N322, the N90 glycan contact pattern displays glycosylation scheme dependence. The ACE2 glycan at N90 interacts with similar RBD residues for both coronaviruses with glycosylation scheme #2 (Fig. S4b). However, with glycosylation scheme #1, the ACE2 glycan at N90 makes more contacts with the SARS-CoV RBD (Fig. 2b). The glycan at N90 differs (in number) by one sugar between the two glycosylation schemes adopted in this study (Fig. S1). The presence of the charged sialic acid (scheme #1) in the glycan at N90 may influence contact between this glycan and RBD. In general, the interacting RBD residues are mostly outside the RBM with the exception of Y491 in SARS-CoV (Tables S1 and S2). However, the larger ACE2 glycan (scheme #1) at N90 makes fewer contacts with the SARS-CoV-2 RBD, and three contacting RBD residues are in the RBM (Table S1). The contact poses and interacting RBD residues for the ACE2 glycan at N90 are found to be similar (scheme #2) between both coronaviruses as shown in Fig. 3. Direct RBD contact of the glycan at N90, which is similar for both coronaviruses with glycan scheme #2, can strengthen overall ACE2-RBD binding (Fig. S4b). However, the difference in contacts between the ACE2 glycan at N90 and RBD with glycosylation scheme #1 indicates that overall ACE2-RBD interactions of SARS-CoV-2 can be impacted by the nature of the ACE2 glycan at N90.

**Figure 3.**
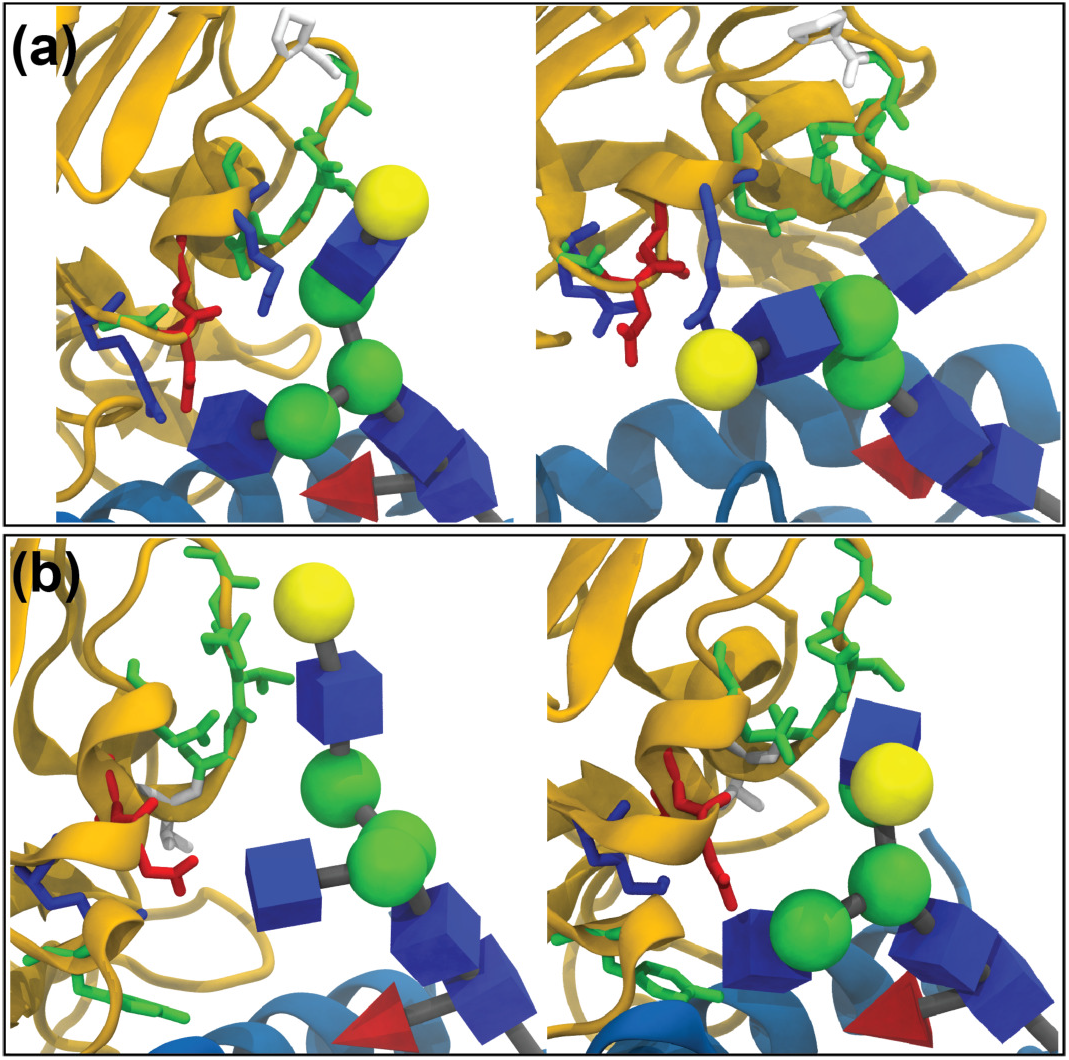
Sample poses for interactions between the ACE2 glycan at N90 and RBD residues of (a) SARS-CoV-2 and (b) SARS-CoV using glycosylation scheme #2 described in Fig. S1. Average number of contacts values are provided in Table S2.

Contact analysis between RBD and ACE2 shows that some ACE2 residues in the ranges E23-N42, L79-Y83, M323-N330, and K353-R357 interacts predominantly with RBDs of both coronaviruses (Fig. S5a-c). Average number of contacts for interacting residues in these ranges are shown in Table S3. Therefore, similar to the glycan-shielding effect reported for the spike protein itself,^13^ ACE2 glycans interacting with the aforementioned ACE2 residues suppress RBD-ACE2 binding and, consequently, may provide protection against coronavirus infections. Using MD simulation of ACE2 without RBD, Mehdipour and Hummer demonstrated that the ACE2 glycan at N90 provides the largest surface coverage compared to other ACE2 glycans.^16^ In the presence of RBDs, we observe that the glycan at N90 interacts with ACE2 residues in a region that overlaps with RBD-ACE2 interactions (residues E23-N42) for both coronaviruses. Average number ACE2 contacts of the ACE2 glycan at N90 for both coronaviruses are provided in Table S4. Note that the ACE2 glycan at N90 makes more contact with ACE2 at the ACE2-RBD interfaces of the SARS-CoV RBD compared to the SARS-CoV-2 RBD. Therefore, the proposed role^16,24^ of the ACE2 glycan at N90 for protection against viral infection is further supported. More direct evidence of the protective role of the ACE2 glycan at N90 has been provided by point mutations of N90 and/or T92. With the exception of T92S, these mutations eliminate the N90 glycosylation. Elimination of the glycosylation at N90 was shown to reduce binding of SARS-CoV-2.^24^ Furthermore, replacing human ACE2 residues 90 to 93 by civet ACE2 residues leads to NLTV → DAKI mutations, also eliminating the N90 glycosylation site. Experimental results demonstrated that the binding affinity of SARS-CoV is reduced because of these mutations.^26^ The net effect of the ACE2 glycan at N90 on coronavirus-ACE2 binding is the combination of these two opposing effects. On the one hand N90 provides protection via occluding the ACE2 residues in the binding interface and on the other it enhances binding by direct interaction with the RBD. The net effect is thus not only coronavirus specific, but also glycosylation scheme dependent.

In conclusion, we determined the roles of ACE2 glycans at N90 and N322 and demonstrated that the ACE2 glycan at N322 contributes to the higher binding affinity of SARS-CoV-2 RBD to ACE2. The ACE2 glycan at N322 facilitates the binding of ACE2 with SARS-CoV-2, while it is occluded by the RBD glycan at N357 in SARS-CoV (N370 in SARS-CoV-2). Therefore, the loss of the glycan at N370 in the RBD of SARS-CoV-2 may be part of its evolution towards stronger virus-host binding. Collectively, we elucidated the roles of different ACE2 glycans in RBD binding and highlighted the differences in glycan-RBD interactions between two coronaviruses, SARS-CoV and SARS-CoV-2. Such differences assist in unraveling the critical role of the glycans the structural and functional differences between SARS-CoV and SARS-CoV-2.

## Acknowledgements

Computational resources were provided through the Extreme Science and Engineering Discovery Environment (XSEDE; TG-MCB130173), which is supported by the National Science Foundation (NSF; ACI-1548562). This work also used the Hive cluster, which is supported by the National Science Foundation under grant number 1828187 and managed by the Partnership for an Advanced Computing Environment (PACE) at the Georgia Institute of Technology.

