## Supplementary Information for "ACE2 glycans preferentially interact with the RBD of SARS-CoV-2 over SARS-CoV"

*School of Physics, Georgia Institute of Technology, Atlanta GA 30332, United States*

### System building, MD simulations, and contact calculations

ACE2 forms a dimeric complex with a transmembrane amino acid transporter, $B^0AT1$. Each peptidase domain (PD) of ACE2 in the dimeric complex can bind to one RBD of SARS-CoV-2.[S1] Furthermore, recent simulations demonstrated that the presence of $B^0AT1$ does not affect the RBD-ACE2 interactions.[S2] Therefore, we only simulated a complex with one coronavirus RBD and one ACE2 peptidase domain in the absence of $B^0AT1$. We performed 2–$\mu$s molecular dynamics (MD) simulations of the RBD-ACE2 complex starting from an x-ray crystal structure of SARS-CoV (PDB ID: 2AJF)[S3] and a cryo-EM structure of SARS-CoV-2 (PDB ID: 6M17);[S1,S4] both had resolutions of 2.9 Å. In order to construct the system with SARS-CoV RBD bound to ACE2, we used chains E and A from the X-ray structure, respectively. There were missing residues in a loop of the SARS-CoV RBD (376-381), which were modeled and added in. The following disulfide bonds between ACE2 residues were added: Cys133 and Cys141, Cys344 and Cys361, and Cys530 and Cys542. For SARS-CoV RBD disulfide bonds were added for Cys323 and Cys348, Cys366 and Cys419, and Cys467 and Cys474. Based on its local electrostatic environment, we protonated Asp350 of ACE2.

For the structure of SARS-CoV-2 RBD bound to ACE2 we used a refined version of chains E and B, respectively, from the cryo-EM structure. All of the disulfide bonds present in the SARS-CoV system were also added here. In addition, disulfide bonds were added for RBD residues Cys336 and Cys361, Cys379 and Cys432, and Cys480 and Cys488. We chose to protonate Asp350 of ACE2 here as well. In all systems ACE2 has a $Zn^{2+}$ ion coordinated to His374, His378, Glu402, and a water molecule in a tetrahedral arrangement. The ion was covalently bound to these residues in our simulations and the force field parameters for the ion complex are described in our previous work.[S5] We retained all crystallographic water molecules. Missing hydrogen atoms were added to all systems, after which they were solvated in a box containing ~64,000 water molecules. We also added $Na^+$ and $Cl^-$ ions to achieve a salt concentration of 150 mM. Glycans were added to each site using the GLYCAM Web server developed by the Woods group (http://glycam.org). Established symbol nomenclatures and 3D representations were consistently used to represent the glycans.[S6,S7]

We used the CHARMM36m force field for proteins,[S8] CHARMM force field for glycans,[S9] and the TIP3P water model[S10] in all simulations. All systems were equilibrated with NAMD

2.13,[S11] which consisted of multiple steps. In the first step, we restrained the proteins and the glycans (if present) for 1 ns, while equilibrating water and ions. In the second step, we restrained only the protein backbones, whereas the side chains and the glycans were equilibrated for 4 ns. In the final equilibration step, all restraints were removed for 10 ns. In the NAMD simulations, constant temperature and pressure were kept at 310 K and 1 atm, respectively, using a Langevin thermostat and piston. The equations of motion were integrated with a 2-fs time step, and long-range electrostatics were evaluated every other time step with particle-mesh Ewald method.[S12] We used a 12-Å cutoff for Lennard-Jones interaction, and a switching function was applied beginning at 10 Å, ensuring a smooth decay to 0. After each of the six systems (two glycosylation schemes as well as no glycosylation for each of SARS-CoV and SARS-CoV-2 RBDs) was equilibrated in NAMD, they were simulated in Amber16. In the Amber16 simulations, we used hydrogen mass repartitioning,[S13,S14] allowing for a time step of 4 fs, and the Monte Carlo barostat for pressure control. All other simulation parameters were identical to those used in the NAMD simulations. For each glycosylation scheme, three replicas were run, while for the systems lacking glycosylation, two replicas were run, giving 32 $\mu$s of simulations in total ([2 glycosylation schemes $\times$ 2 systems $\times$ 3 replicas $\times$ 2 $\mu$s] + [2 systems $\times$ 2 replicas $\times$ 2 $\mu$s of non-glycosylated simulations]).

Contact calculations were performed using a cutoff distance of 3.5 Å. Therefore, when two heavy atoms from two different selections come with 3.5 Å, we count that as one contact. To calculate all average properties we pooled data from all independent MD simulations.

### Data availability

Simulation trajectories are available at the NSF MolSSI COVID-19 Molecular Structure and Therapeutics Hub at https://covid.molssi.org.

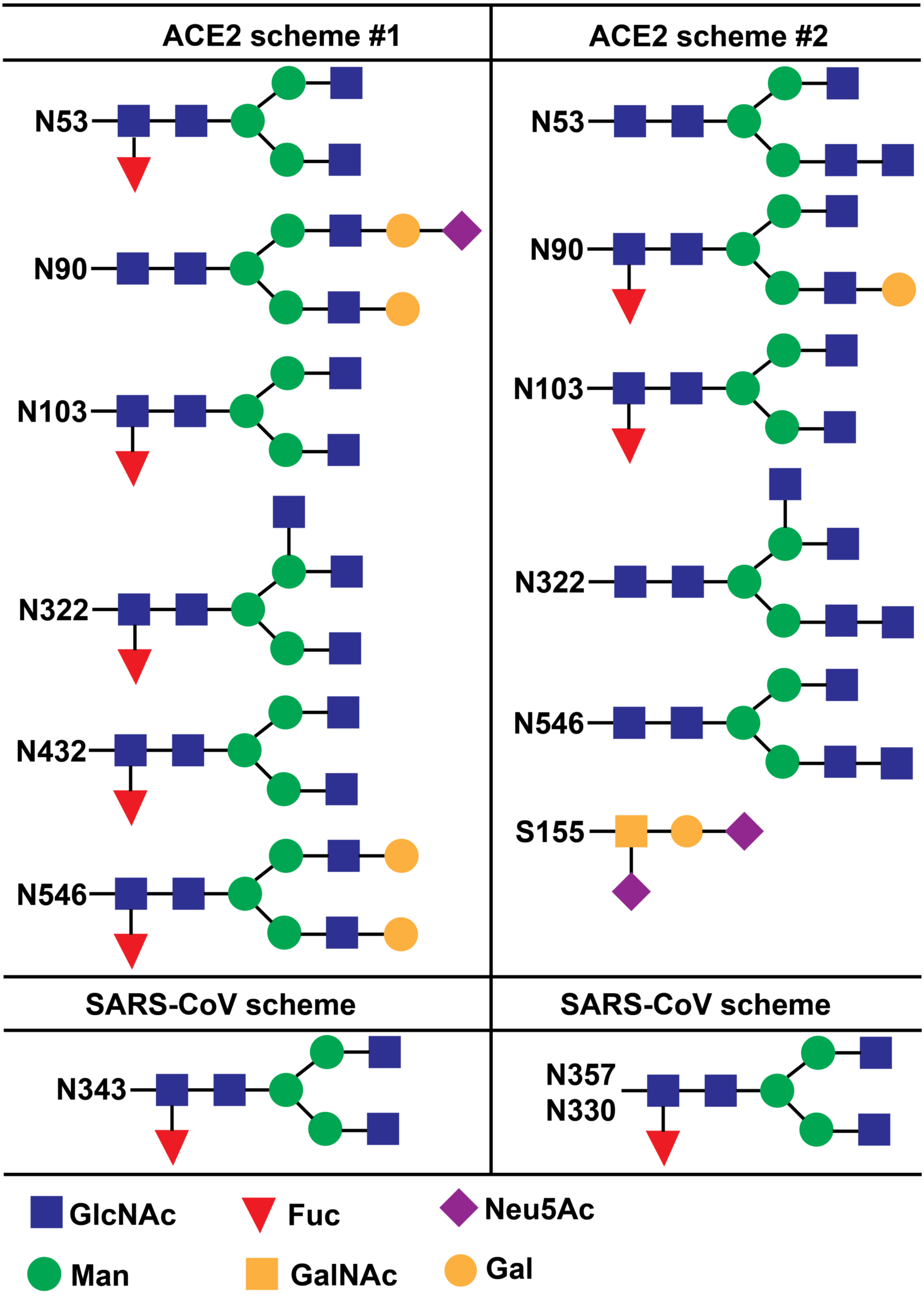


Figure S1: Glycosylation schemes adopted in this study following populations proposed by Zhao et al.[S15] and Shajahan et al. [S16]

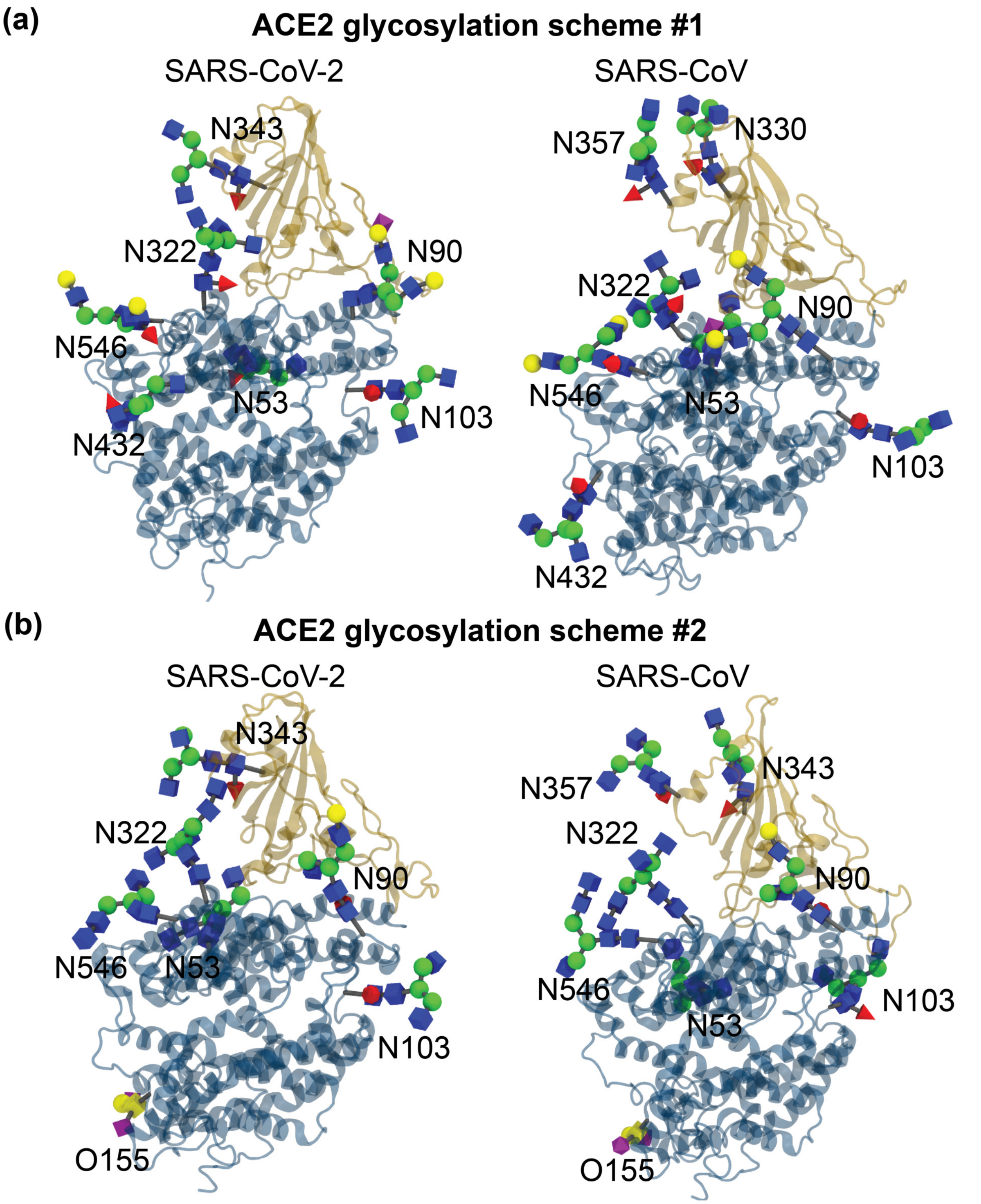


Figure S2: Fully glycosylated model systems of RBD-ACE2 complexes for two coronaviruses with (a) scheme #1 and (b) scheme #2

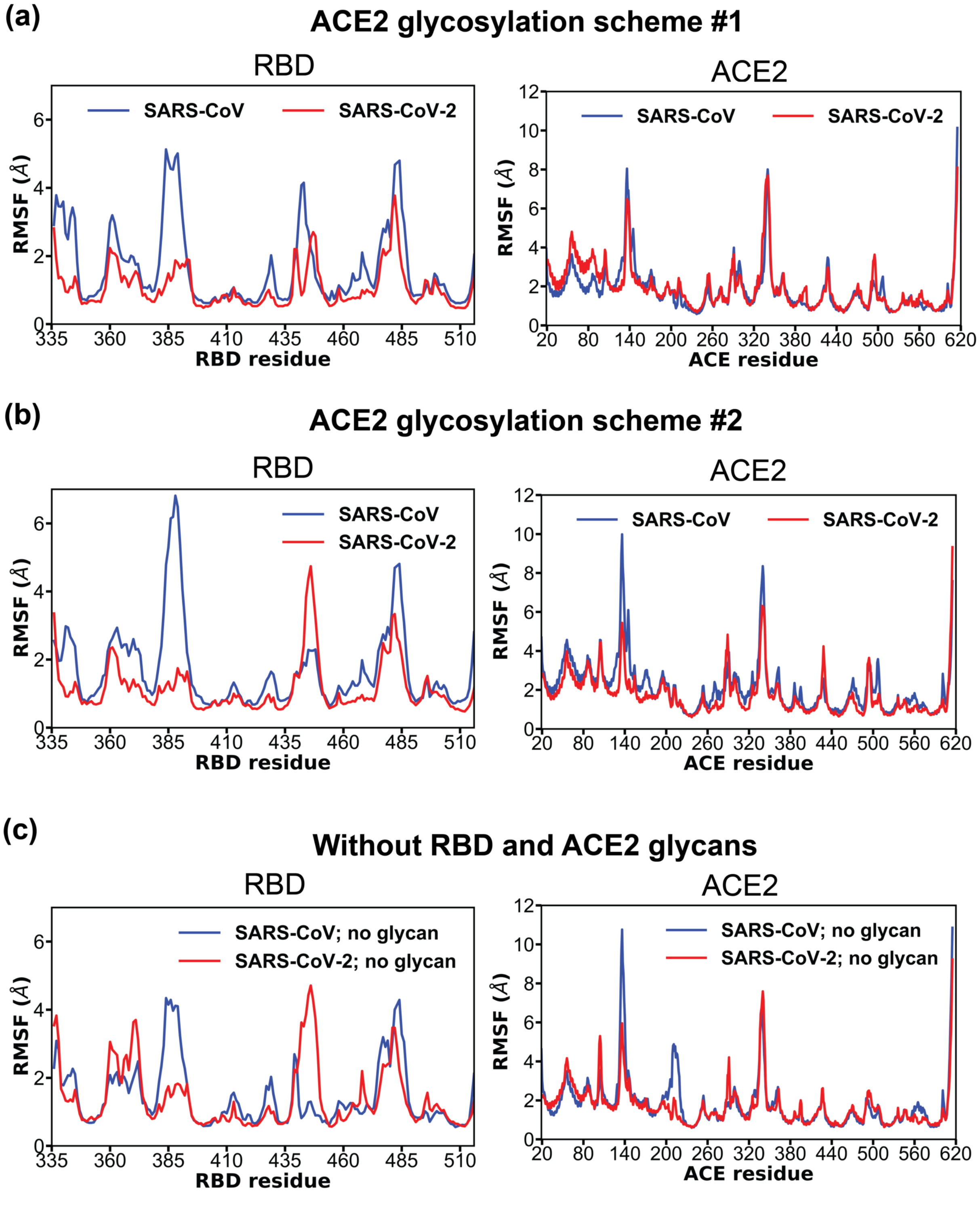


Figure S3: Root mean square fluctuation (RMSF) of coronavirus RBDs and ACE2 in each RBD-ACE2 complex with glycosylation (a) scheme #1, (b) scheme #2, and (c) without glycans. Glycosylations schemes are defined in Figure S1. Simulation data from multiple independent 2-$\mu$s simulations (with glycans: $3 \times 2$ $\mu$s, without glycans: $2 \times 2$ $\mu$s) were pooled for RMSF calculations.

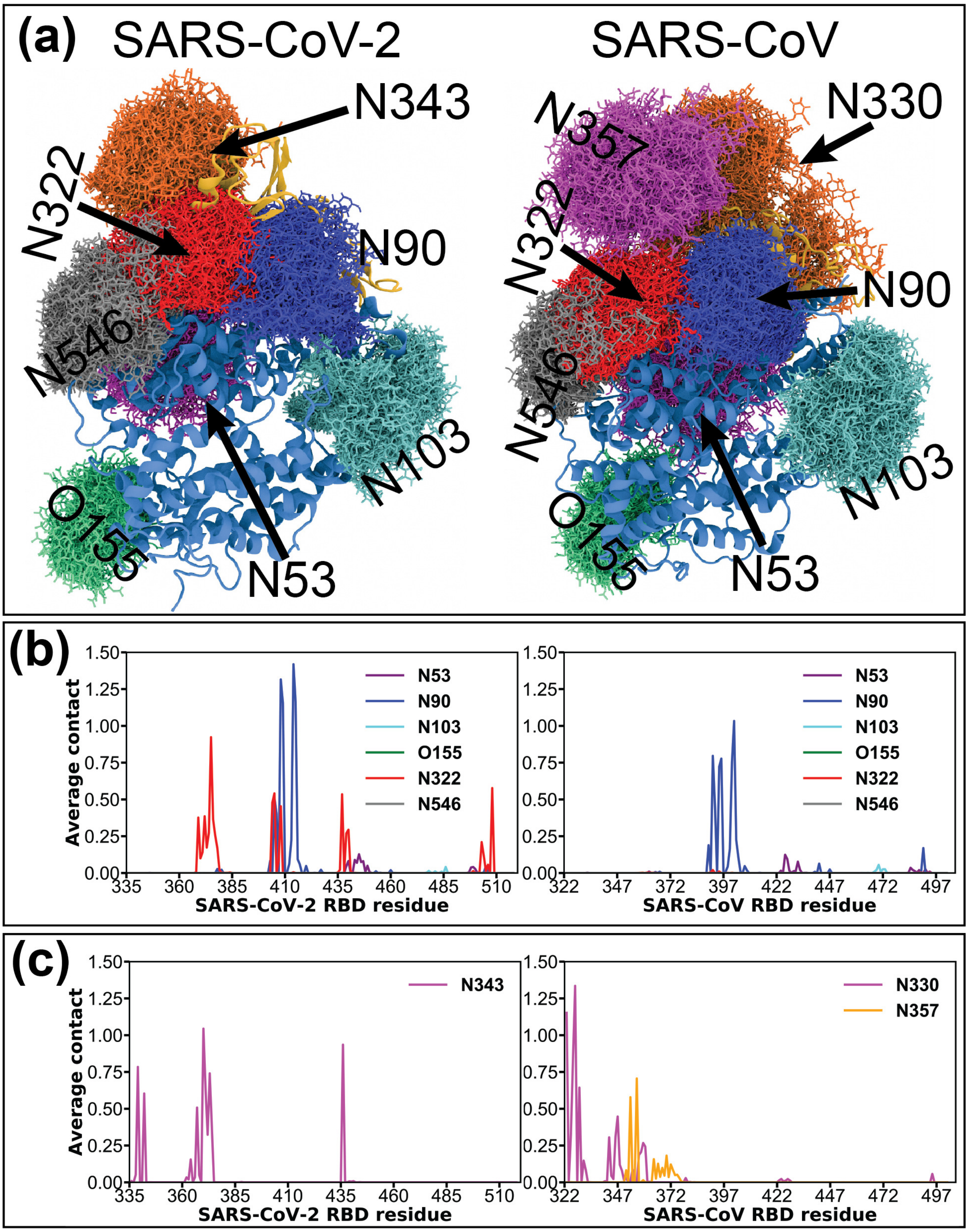


Figure S4: (a) Comparison of glycan coverage (b) average contact between ACE2 glycans and RBD residues, and (c) average contact between RBD glycans and RBD residues of SARS-CoV-2 (left) and SARS-CoV (right) with glycosylation scheme #2 defined in Figure S1.

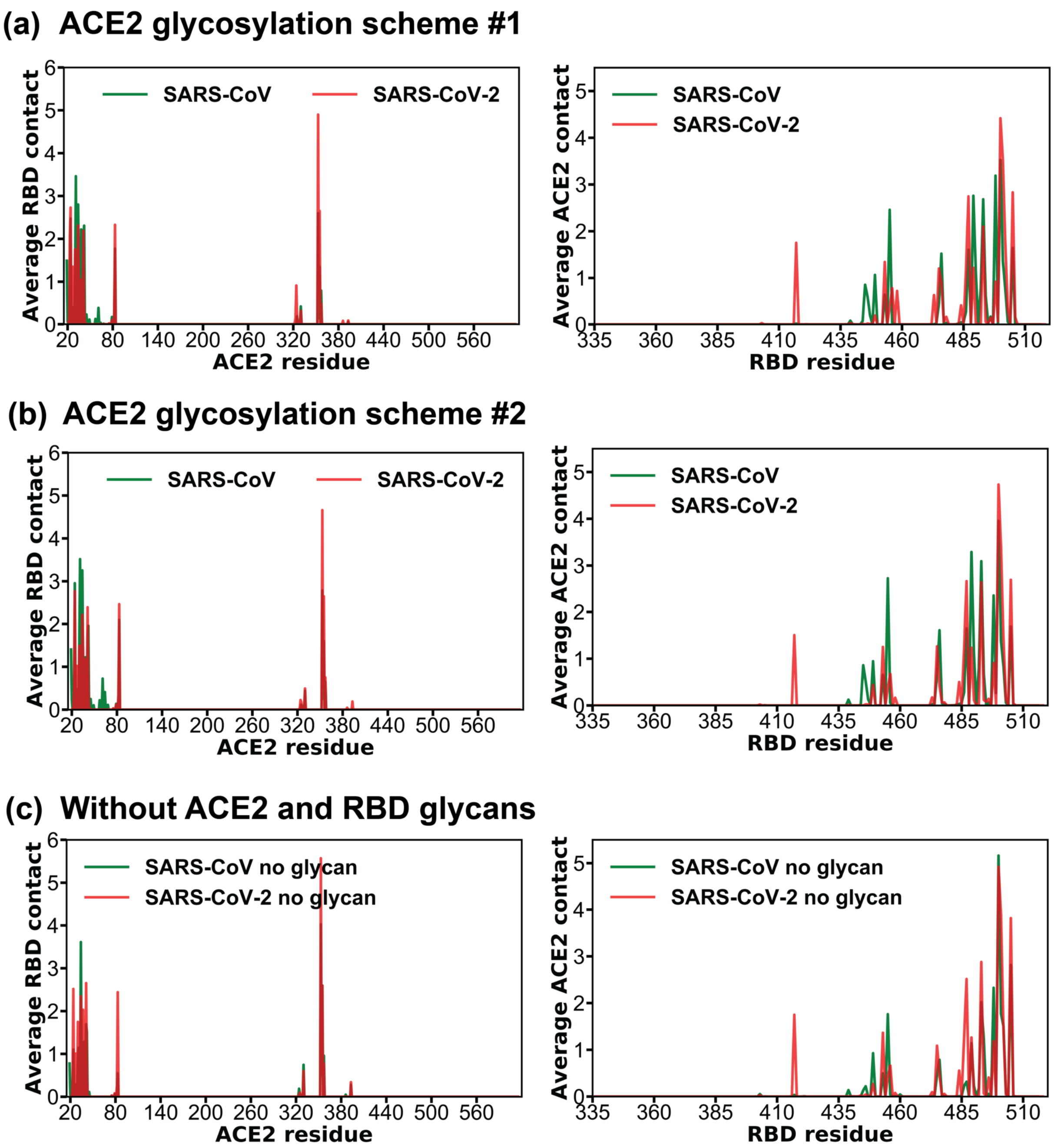


Figure S5: Average contact between RBD and ACE2 residues with glycosylation (a) scheme #1, (b) scheme #2, and (c) without glycans. Glycosylations schemes are defined in Figure S1. Simulation data from multiple independent 2-$\mu$s simulations (with glycans: 3 $\times$ 2 $\mu$s, without glycans: 2 $\times$ 2 $\mu$s) were pooled for this analysis.

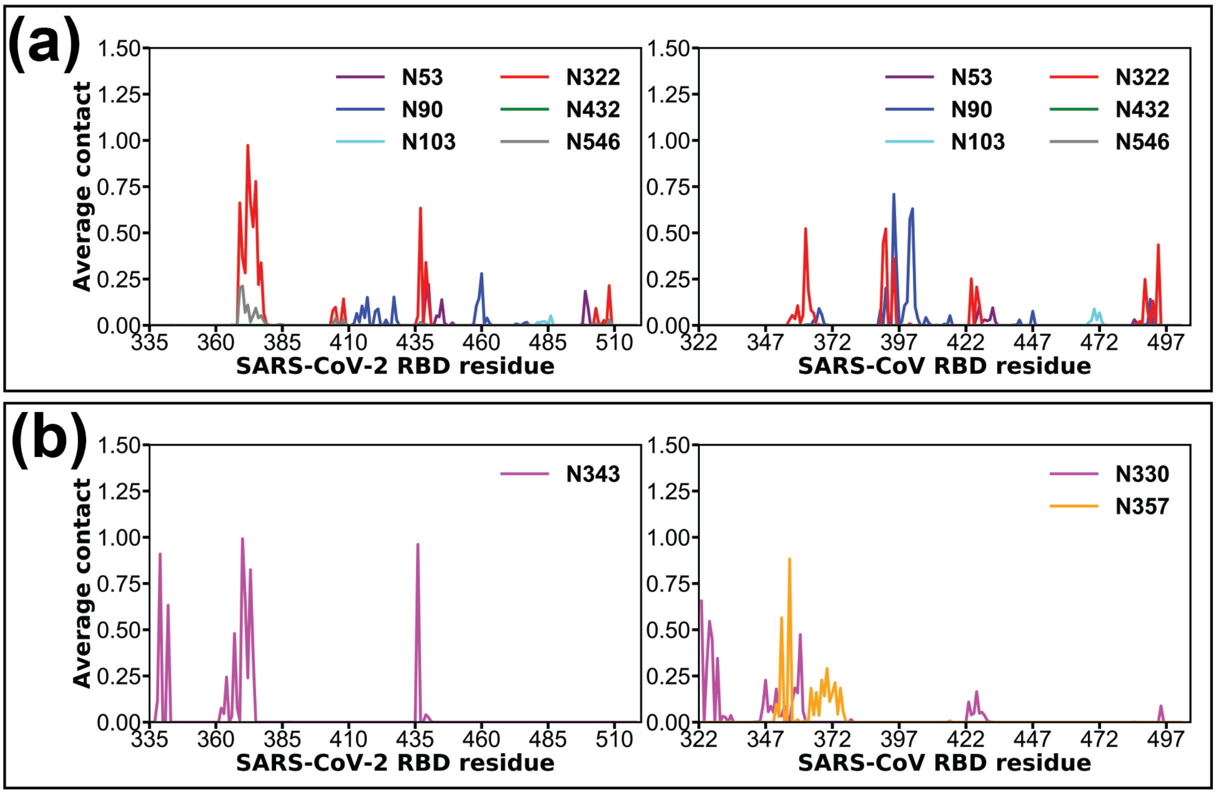


Figure S6: (a) Average contact between all ACE2 glycans and RBD residues, and (c) average contact between RBD glycans and RBD residues of SARS-CoV-2 (left) and SARS-CoV (right) with glycosylation scheme #1 defined in Figure S1. Results with glycosylation scheme #2 is presented in Figures S4b,c.

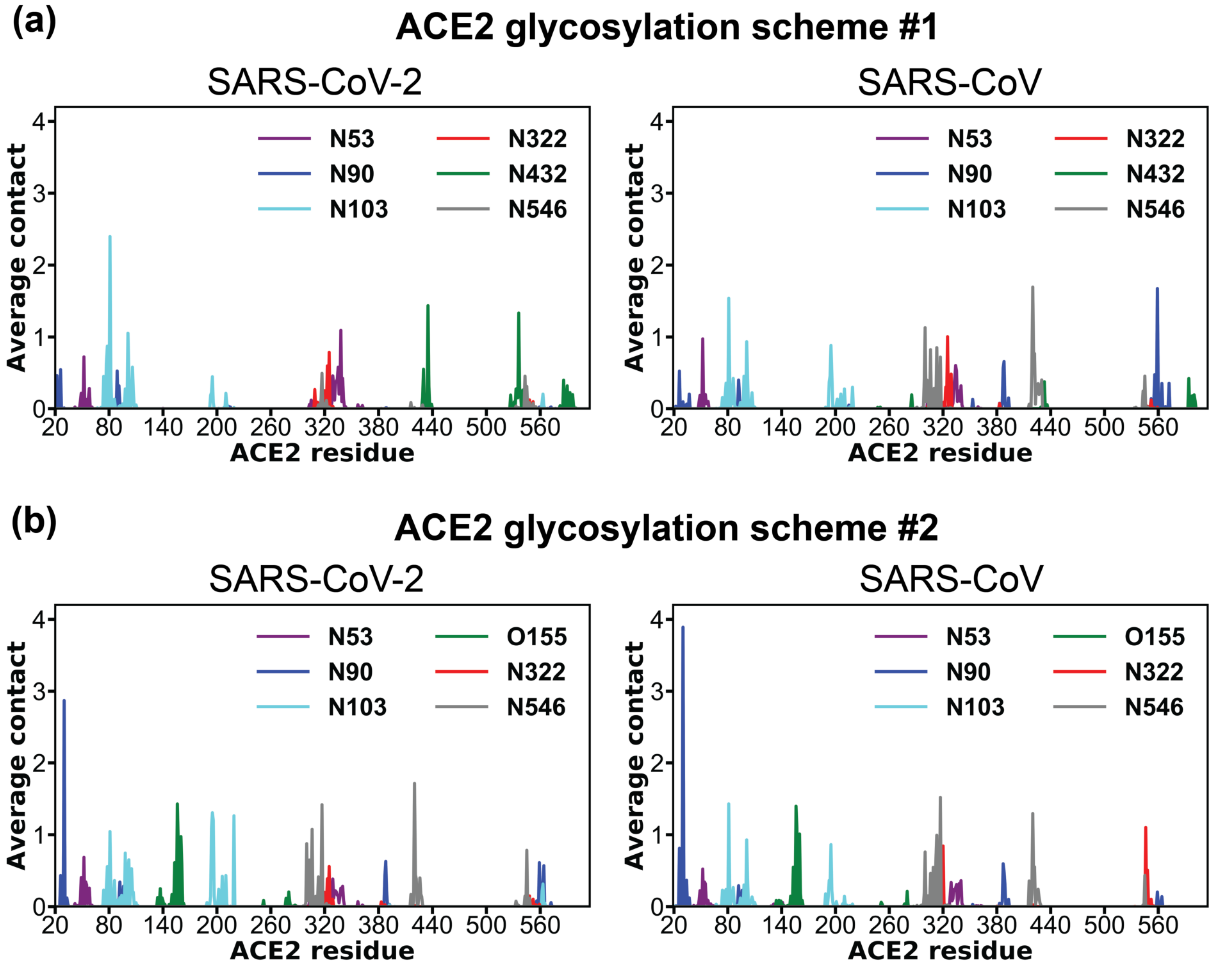


Figure S7: Average contact between ACE2 glycan and ACE2 protein residues with glycosylation (a) scheme #1 and (b) scheme #2. Glycosylations schemes are defined in Figure S1. Simulation data from three independent 2-$\mu$s simulations were pooled for this analysis.

Table S1: Average contact of ACE2 glycans with two coronavirus RBDs for glycan scheme #1 defined in Figure S1. Average contact values >0.1 (obtained from 3 $\times$ 2-$\mu$s simulations) are tabulated. Non-conserved residues are highlighted in bold font.

| ACE2 glycan | **Glycan scheme #1** | | | |
|---|---|---|---|---|
| | SARS-CoV-2 residue | <contact> | SARS-CoV residue | <contact> |
| **N90** | Asp405 | _ | Asp392 | 0.20 |
| | Arg408 | _ | Arg395 | 0.71 |
| | Gln409 | _ | Gln396 | 0.32 |
| | Pro412 | _ | Pro399 | 0.11 |
| | Gly413 | _ | Gly400 | 0.13 |
| | Gln414 | _ | Gln401 | 0.57 |
| | Thr415 | 0.10 | Thr402 | 0.63 |
| | Gly416 | _ | Gly403 | 0.10 |
| | **Lys417** | 0.15 | **Val404** | _ |
| | Asp427 | 0.15 | Asp414 | _ |
| | Lys458 | 0.10 | His445 | _ |
| | **Ser459** | 0.15 | **Gly446** | _ |
| | **Asn460** | 0.28 | **Lys447** | _ |
| | Tyr505 | _ | Tyr491 | 0.14 |
| **N322** | Tyr369 | 0.66 | Tyr356 | _ |
| | Asn370 | 0.36 | Asn357 | _ |
| | Ser371 | 0.28 | Ser358 | _ |
| | **Ala372** | 0.97 | **Thr359** | _ |
| | **Ser373** | 0.66 | **Phe360** | _ |
| | Phe374 | 0.53 | Phe361 | _ |
| | Ser375 | 0.78 | Ser362 | 0.52 |
| | Thr376 | 0.22 | Thr363 | 0.19 |
| | Phe377 | 0.34 | Phe364 | _ |
| | Gly404 | _ | Gly391 | 0.45 |
| | Asp405 | 0.10 | Asp392 | 0.52 |
| | Arg408 | 0.14 | Arg395 | 0.36 |
| | Asn437 | 0.63 | Asn424 | 0.25 |
| | **Asn439** | 0.34 | **Arg426** | 0.21 |
| | Asn440 | 0.14 | Asn427 | 0.11 |
| | **Val503** | _ | **Ile489** | 0.25 |
| | Gln506 | _ | Gln492 | 0.13 |
| | Tyr508 | 0.21 | Tyr494 | 0.44 |
| **N546** | Tyr369 | 0.20 | Tyr356 | _ |
| | Asn370 | 0.21 | Asn357 | _ |
| | **Ala372** | 0.11 | **Thr359** | _ |
| **N53** | **Asn439** | 0.17 | **Arg426** | _ |
| | Asn440 | 0.22 | Asn427 | _ |
| | **Val445** | 0.14 | **Ser432** | _ |
| | **Pro499** | 0.18 | **Thr485** | _ |
| | Thr500 | 0.10 | Thr486 | _ |

Table S2: Average contact of ACE2 glycans with two coronavirus RBDs for glycan scheme #2 defined in Figure S1. Average contact values >0.1 (obtained from 3 × 2-$\mu$s simulations) are tabulated. Non-conserved residues are highlighted in bold font.

| ACE2 glycan | **Glycan scheme #2** | | | |
|---|---|---|---|---|
| | SARS-CoV-2 residue | <contact> | SARS-CoV residue | <contact> |
| **N90** | **Arg403** | – | **Lys390** | 0.19 |
| | Asp405 | 0.51 | Asp392 | 0.80 |
| | **Glu406** | 0.40 | **Asp393** | 0.46 |
| | Arg408 | 1.32 | Arg395 | 0.72 |
| | Gln409 | 1.16 | Gln396 | 0.78 |
| | Gly413 | 0.22 | Gly400 | 0.16 |
| | Gln414 | 1.42 | Gln401 | 0.75 |
| | Thr415 | 1.16 | Thr402 | 1.03 |
| | | | Gly403 | 0.23 |
| | | | Val404 | 0.11 |
| | | | Tyr491 | 0.17 |
| **N322** | Tyr369 | 0.38 | Tyr356 | – |
| | Asn370 | 0.10 | Asn357 | – |
| | Ser371 | 0.14 | Ser358 | – |
| | **Ala372** | 0.39 | **Thr359** | – |
| | **Ser373** | 0.17 | **Phe360** | – |
| | Phe374 | 0.27 | Phe361 | – |
| | Ser375 | 0.92 | Ser362 | – |
| | Thr376 | 0.36 | Thr363 | – |
| | Phe377 | 0.25 | Phe364 | – |
| | Lys378 | 0.17 | Lys365 | – |
| | Gly404 | 0.48 | Gly391 | – |
| | Asp405 | 0.54 | Asp392 | – |
| | Arg408 | 0.45 | Arg395 | – |
| | Asn437 | 0.53 | Asn424 | – |
| | **Asn439** | 0.27 | **Arg426** | – |
| | Asn440 | 0.29 | Asn427 | – |
| | **Val503** | 0.21 | **Ile489** | – |
| | Gly504 | 0.12 | Gly490 | – |
| | Tyr508 | 0.58 | Tyr494 | – |

Table S3: Average RBD-ACE2 contact calculated for two glycosylations schemes defined in Figure S1. Simulation data from multiple independent 2-$\mu$s simulations (with glycans: 3 $\times$ 2 $\mu$s, without glycans: 2 $\times$ 2 $\mu$s) were pooled for this analysis.

| ACE2 residue | SARS-CoV-2 | | | SARS-CoV | | |
|---|---|---|---|---|---|---|
| | scheme #1 | scheme #2 | no glycan | scheme #1 | scheme #2 | no glycan |
| Glu23 | 2.1 | 0.5 | 0.3 | | | |
| Gln24 | 2.7 | 2.8 | 2.5 | 2.5 | 3.0 | 1.1 |
| Thr27 | 1.3 | 1.0 | 1.0 | 0.4 | 0.5 | 0.3 |
| Phe28 | 0.7 | 0.7 | 0.8 | 0.3 | 0.2 | 0.2 |
| Asp30 | 1.8 | 1.5 | 1.7 | 1.2 | 1.6 | 0.8 |
| Lys31 | 1.2 | 1.5 | 1.4 | 3.5 | 3.5 | 1.1 |
| His34 | 2.3 | 2.2 | 2.3 | 2.8 | 3.3 | 3.6 |
| Glu35 | 0.9 | 1.3 | 1.6 | 0.7 | 0.6 | 0.4 |
| Glu37 | 1.0 | 1.2 | 2.0 | 0.5 | 0.4 | 1.1 |
| Asp38 | 0.2 | 0.4 | 0.4 | 2.2 | 1.2 | 1.3 |
| Tyr41 | 2.2 | 2.4 | 2.7 | 2.1 | 1.8 | 1.7 |
| Gln42 | 0.4 | 0.4 | 0.5 | 2.3 | 2.0 | 1.5 |
| Leu79 | 0.1 | 0.1 | 0.1 | 0.2 | 0.1 | 0.1 |
| Met82 | 0.2 | 0.3 | 0.3 | 0.1 | 0.2 | 0.0 |
| Tyr83 | 2.3 | 2.5 | 2.4 | 1.8 | 2.1 | 0.5 |
| Met323 | 0.1 | | | | | |
| Thr324 | 0.9 | 0.2 | | | | 0.2 |
| Gln325 | 0.2 | 0.2 | 0.1 | 0.2 | 0.1 | 0.1 |
| Glu329 | | | | | 0.1 | 0.1 |
| Asn330 | 0.3 | 0.5 | 0.6 | 0.4 | 0.4 | 0.7 |
| Lys353 | 4.9 | 4.7 | 5.6 | 2.6 | 2.8 | 4.0 |
| Gly354 | 0.8 | 0.5 | 0.5 | 0.1 | 0.3 | 0.5 |
| Asp355 | 2.7 | 2.6 | 2.6 | 1.3 | 1.6 | 2.6 |
| Arg357 | 0.5 | 0.8 | 0.8 | 0.8 | 0.6 | 1.0 |

Table S4: Average N90-ACE2 contact (in the RBD-ACE2 interface) calculated for two glycosylations schemes defined in Figure S1. Simulation data from three independent 2-$\mu$s simulations were pooled for this analysis.

| ACE2 residue | SARS-CoV-2 | | SARS-CoV | |
|---|---|---|---|---|
| | scheme #1 | scheme #2 | scheme #1 | scheme #2 |
| Glu22 | 0.5 | | | |
| Lys26 | 0.5 | 0.4 | 0.5 | 0.8 |
| Leu29 | | 0.2 | | 0.2 |
| Asp30 | | 2.9 | 0.1 | 3.9 |
| Asn33 | | 0.1 | | 0.3 |
| His34 | | | | 0.4 |
| Glu37 | | | 0.2 | 0.1 |
